# Evidence of Disrupted-in Schizophrenia 1 (DISC1) as an arsenic binding protein and implications regarding its role as a translational activator

**DOI:** 10.1101/2023.06.14.544995

**Authors:** Muneaki Watanabe, Tung Mei Khu, Grant Warren, Juyoung Shin, Charles E. Stewart, Julien Roche

## Abstract

Disrupted-in-schizophrenia-1 (DISC1) is a scaffold protein that plays a pivotal role in orchestrating signaling pathways involved in neurodevelopment, neural migration, and synaptogenesis. Among those, it has recently been reported that the role DISC1 in the Akt/mTOR pathway can shift from a global translational repressor to a translational activator in response to oxidative stress induced by arsenic. In this study we are providing evidence that DISC1 can directly bind arsenic via a C-terminal cysteine motif (C-X-C-X-C). A series of fluorescence-based binding assays were conducted with a truncated C-terminal domain construct of DISC1 and a of series of single, double, and triple cysteine mutants. We found that arsenous acid, a trivalent arsenic derivative, specifically binds to the C-terminal cysteine motif of DISC1 with low micromolar affinity. All three cysteines of the motif are required for high-affinity binding. Electron microscopy experiments combined with in silico structural predictions revealed that that the C-terminal of DISC1 forms an elongated tetrameric complex. The cysteine motif is consistently predicted to be located within a loop, fully exposed to solvent, providing a simple molecular framework to explain the high-affinity of DISC1 toward arsenous acid. This study sheds light on a novel functional facet of DISC1 as an arsenic binding protein and highlights its potential role as both a sensor and translational modulator within the Akt/mTOR pathway.

## INTRODUCTION

*Disrupted-in-schizophrenia-1* (*DISC1*) was first identified at the breakpoint of a balanced chromosomal translocation (1;11) (q42.1;q14.3) in a Scottish family with a history of schizophrenia and related psychiatric disorders.^1,2^ Subsequent studies have shown that several genetic variations of *DISC1* are linked to increased risks of mental disorders and neuronal diseases.^3-5^ The product of *DISC1* is an 854 amino acid-long protein that serves as a scaffold hub at the center stage of major neuronal signaling pathways.^6^ A growing wealth of information on the physiological role of DISC1 describes its importance in cellular functions such as proliferation, neuronal development, and synaptogenesis.^7-10^ Notably, accumulating evidence supports the participation of DISC1 in translational processes in association with the Akt/mTOR signaling pathway.^11,12^ It has for instance been reported that DISC1 directly binds and inhibits GSK3β, which is itself inhibited by Akt.^13^ DISC1 is known to interact with the actin-binding protein girdin (also known as KIAA1212), a key activator of the Akt/mTOR pathway.^11,12^ DISC1 has also been reported to directly associate with polyribosomes as a translational activator.^14^ Furthermore, a recent study by Fuentes-Villalobos et al. has highlighted the role of DISC1 in maintaining homeostatic control of protein synthesis during oxidative stress.^15^ In non-stress conditions DISC1 acts as an overall inhibitor of the Akt/mTOR pathway by sequestering the activator girdin, therefore exerting a negative control on the translation processes associated with this pathway (**Fig. 1A**).^11-12^ Interestingly, the role of DISC1 as a translational inhibitor is reversed when cells are treated with sodium arsenite, inducing oxidative stress.^15^ Indeed, in oxidative stress conditions, DISC1 appears to act as a general enhancer of protein synthesis by interacting with eiF3, a key component of the translational machinery that recruits the small ribosomal subunit 40S (**Fig. 1B**).^14,15^ The study by Fuentes-Villalobos et al. therefore suggests that DISC1 is required for rescuing protein synthesis under oxidative stress conditions.^15^ Yet the molecular mechanisms underlying the switch of DISC1 function from a general inhibitor to an activator of protein synthesis remain elusive.

**Figure 1.**
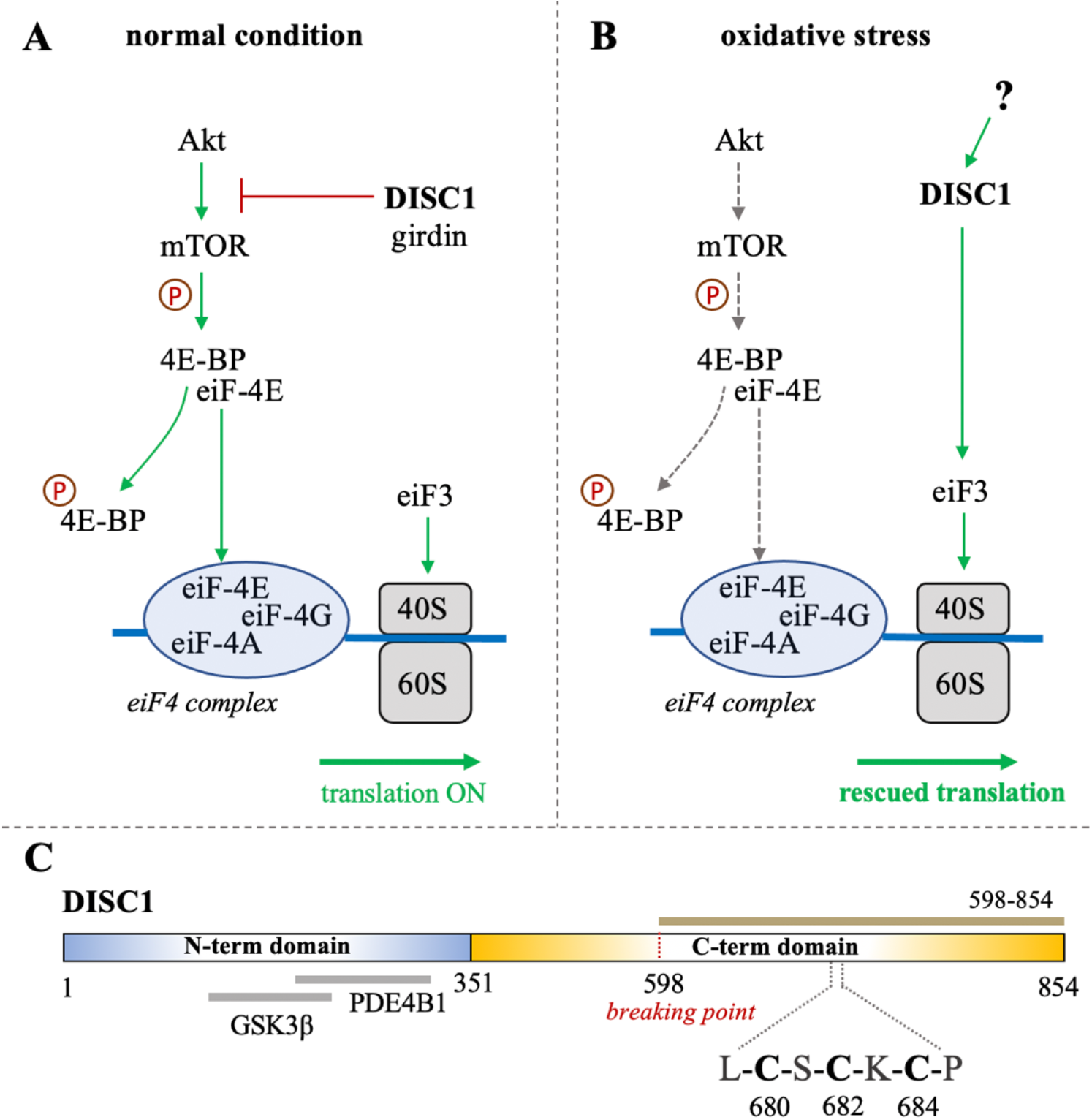
**(A-B)** Schematic representation of the role of DISC1 as translational modulator within the Akt/mTOR pathway, in (**A**) normal conditions and in (**B**) in situation of oxidative stress. In normal conditions, DISC1 indirectly modulates the translational activity of the eiF4 complex by inhibiting the Akt-mediated activation of mTOR through sequestration of the actin-binding protein girdin. In oxidative stress conditions, the Akt/mTOR pathway is globally inhibited but DISC1 can rescue translational activity by activating eiF3. (**C**) Domain organization of DISC1 depicting the predominantly disordered N-terminal domain (residues 1-351) in blue that harbors known binding sites to GSK3β and PDE4B1. The C-terminal domain predicted to be rich in helical and coil motifs and containing the cysteine motif composed of C680, C682, and C684, is shown in yellow. The C-terminal construct used in this study (598-854) is highlighted in brown.

Bioinformatics analysis suggest that DISC1 is composed of a predominantly disordered N-terminal domain (residues 1-350) and a helical C-terminal domain (residues 351-854).^16^ The N-terminal domain contains two nuclear-localization signals as well as the binding sites to key protein partners, including the phosphodiesterase PDE4B1 and GSK3β (**Fig. 1C**).^16,17^ As often observed with large scaffold proteins, structural disorder appears to be tightly connected to DISC1 function. Coarse-grained molecular simulations have indeed shown that the conformational ensemble of DISC1 is predominantly shaped by transient “fuzzy” interactions mediated by the disordered regions of the protein.^18^ The C-terminal domain is predicted to be more helical with several coiled-coil regions and helical hairpins known as UVR motifs.^16^ Structural properties of short fragments of the C-terminal domain have been characterized via various biophysical techniques, including NMR spectroscopy and SAXS.^19-21^ The C-terminal domain harbors several mutations that are linked to increased risks of mental illness, including the polymorphism S704C^22^ and a frameshift mutation downstream of L807 that leads to pathological aggregation of DISC1.^23^ Moreover, the balanced chromosomal translocation discovered in the Scottish family results in the truncation of a large C-terminal fragment (residues 598-854) (**Fig. 1C**).

Interestingly, analysis of the C-terminal domain sequence reveals the presence of a cysteine motif (C-X-C-X-C) at position 680-684 (**Fig. 1C**), which is typically found in arsenic-binding proteins. It is indeed well-known that arsenic can bind to the sulfhydryl group of cysteines.^24^ This led us to investigate whether DISC1 can directly bind arsenic derivatives. We hypothesize that the switch of function from a global inhibitor to an activator of protein synthesis in oxidative stress conditions may be triggered by direct binding of arsenic. To test this hypothesis, we designed a fluorescence-based binding assay and measured the binding affinity of the C-terminal domain of DISC1 to arsenic using a series of single, double, and triple cysteine mutants. We found that arsenous acid specifically binds to the C-terminal cysteine motif of DISC1 (C680, C682, and C684) with low micromolar binding affinity. All three cysteines of the motif are required for high-affinity binding. These results highlight the potential role of DISC1 as a as a cellular sensor for toxic metals such as arsenic.

## RESULTS

Characterization of DISC1 at a molecular level has historically been hampered by its low solubility and high aggregation propensity when expressed recombinantly. For this study, we focused on a truncated C-terminal construct of the human DISC1 corresponding to residues 598-854 (**Fig. 1C**). This construct encompasses the structured regions S (residues 635–738) and C (residues 691–836) identified by Korth and coworkers.^25^ Importantly, it also contains the cysteine motif (C680, C682, and C684) that constitutes the main focus of the present study. When co-expressed with an N-terminal hexa-histidine tag, we were able to express and purify the C-terminal construct of DISC1 with high-yield. The purified protein elutes predominantly as a tetrameric species (**Fig. S1**) and shows no sign of aggregation at the concentration employed in our assays (2 μM). For sake of simplicity, this C-terminal construct (598-854) will be referred as “**WT**” throughout the manuscript.

### 1. Arsenous acid binds to the C-terminal region of DISC1

To investigate whether DISC1 could act as a cellular sensor for toxic heavy metals such as arsenic, we designed a fluorescence assay for monitoring potential binding to the C-terminal DISC1 (WT) construct in an in-vitro environment. We observed a significant decrease in tryptophan fluorescence intensity when the WT construct was titrated with trivalent arsenous acid (As(OH)_3_), a reactive species known to bind with high affinity to cysteine thiol groups of arsenic binding proteins. We then compared the binding properties of WT to a series of variants to decipher the role played by the cysteine motif (C680, C682, and C684) in mediating interaction with arsenous acid (**Table 1**). All variants show similar retention time by size exclusion chromatography, indicating that the engineered mutations did not affect the quaternary structure of the DISC1 C-terminal domain. Binding curves were collectively fitted for all variants (including WT) to a simple binding saturation model to extract the specific and non-specific dissociation constants (K_d_ (high) and K_d_ (low) respectively, **Fig. 2**) and maximum binding capacity (**Table 1**).

**Table 1.** List of constructs with corresponding mutations used in this study. The amino acid numbers refer to the human sequence of DISC1 (**Fig. 1C**). Maximum binding capacity reported for each variant was obtained via a collective fit of all the titration profiles to a two-site specific binding model.

| Name | Construct/mutations | Maximum binding capacity (%) |
| --- | --- | --- |
| **WT** | DISC1 598-854 | 95 ± 6 |
| **WT\*** | DISC1 598-854<br>C698S/C762S/C782S/C844S | 75 ± 4 |
| **C680S\*** | DISC1 598-854<br>C680S/C698S/C762S/C782S/C844S | 40 ± 4 |
| **C682S\*** | DISC1 598-854<br>C682S/C698S/C762S/C782S/C844S | 25 ± 4 |
| **C684S\*** | DISC1 598-854<br>C684S/C698S/C762S/C782S/C844S | 24 ± 3 |
| **C680S/C682S\*** | DISC1 598-854<br>C680S/C682S/C698S/C762S/C782S/C844S | 25 ± 5 |
| **C680S/C684S\*** | DISC1 598-854<br>C680S/C684S/C698S/C762S/C782S/C844S | 25 ± 4 |
| **C682S/C684S\*** | DISC1 598-854<br>C682S/C684S/C698S/C762S/C782S/C844S | 35 ± 4 |
| **C680S/C6802S/C684S\*** | DISC1 598-854<br>C680S/C682S/C684S/C698S/C762S/C782S/C844S | 17 ± 4 |

**Figure 2.**
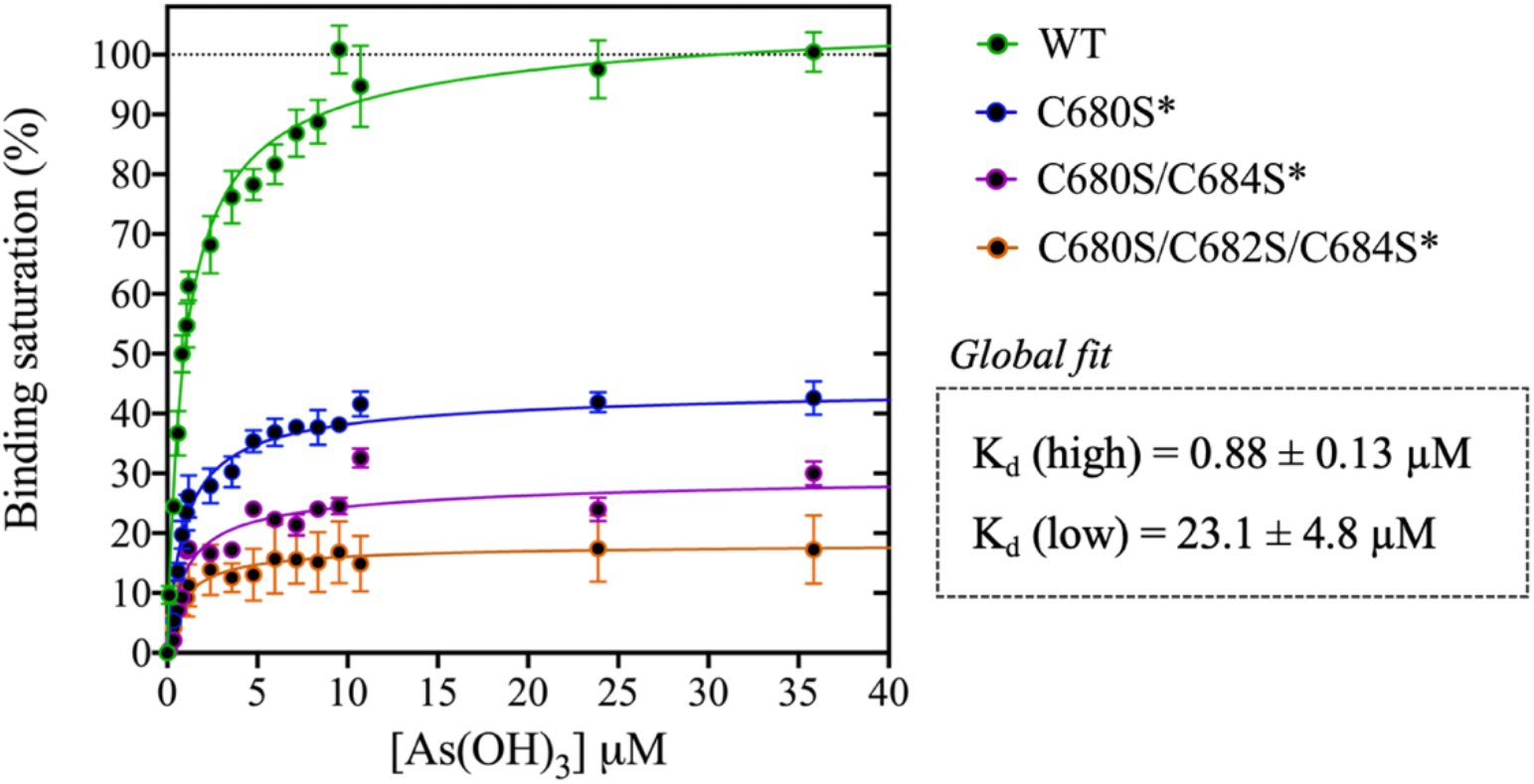
Titration of DISC1 WT construct (598-854) and single (C680S*), double (C680S/C684S*), and triple (C680S/C684S/C682S*) cysteine mutants with arsenous acid. The plot depicts overall binding saturation as a function of arsenous acid concentration derived from fluorescence-based assays. Error bars represents standard deviations from three independent measurements. Lines correspond to a collective fit of all data presented in Table 1 using a simple two-site specific binding model. The dissociation constants calculated from the global fit (K_d_ (high) and K_d_ (low) are shown in insert.

We first examined a variant for which all cysteines, beside C680, C682, and C684, were mutated to serine. This variant named WT* displayed strong binding to arsenous acid, with a maximum binding capacity yet significantly lower than WT (75% compared to 95%, **Table 1, Fig. S2A**). These results suggest that non-specific binding of arsenous acid to cysteines outside the cysteine motif C680, C682, and C684 accounts for nearly 20% of the total binding capacity measured for the WT construct. Next, we consider a series of single, double, and triple mutants engineered in the WT* background (i.e. all cysteines beside C680, C682, and C684 mutated to serine, **Table 1**). All three single-point mutants show drastic reduction in binding capacity indicating that all three cysteines of the cysteines motif are required for optimal binding to trivalent arsenic (**Fig 2** and **Fig. S2B**). Interestingly, we found that C680S* retains a higher binding capacity than C682S* and C684S* (**Table 1**), suggesting a difference in solvent exposure or conformational flexibility among the three cysteines of the motif. The binding capacity of double cysteine-to-serine mutants did not show further decrease in binding capacity, confirming that optimal binding to arsenous acid is not possible without all three cysteines of the motif (**Fig. 2, Fig. S2C**, and **Table 1**). Finally, we examined a variant for which all cysteines have been mutated to serine (C680S/C6802S/C684S*). This variant displays a very low but non-zero binding capacity (**Fig. 2**), suggesting that polar amino acids other than cysteines provide a small but non-negligeable contribution to arsenous acid binding. All together these results indicate that the trivalent arsenous acid binds with high affinity (K_d_ (high) = 0.88 ± 0.13 μM, **Fig. 2**) to a cysteine motif composed of C680S, C682S, and C684S, found in the C-terminal domain of DISC1. We found that all three cysteines within the motif are required to achieve optimal binding to arsenous acid.

### 2. Arsenic binding motif is exposed to solvent in DISC1 tetrameric state

To determine the structural and molecular basis for DISC1 binding to arsenic, we then collected a series of negative-contrast electron micrographs of the “WT” DISC1 C-terminal construct (598-854) in the absence and in the presence of arsenous acid. The micrographs collected in the absence of arsenous acid show two major populations: a heterogeneous ensemble of high-molecular species representing higher oligomers, fibrils, and amorphous aggregates, and a homogeneous population of oligomeric particles of apparent spheroidal shape (**Fig. 3A**). 3D reconstruction of the smaller oligomeric species show a symmetrical spheroid with a major axis of 10-12 nm and shorter axis of 5-6 nm (**Fig. 3C**, upper panel). The dimension of these particles are within with the range of expected dimensions for the tetrameric species observed by size-exclusion chromatography (**Fig. S1**). No significant change in the population distribution was observed upon incubation with arsenous acid, suggesting that arsenic binding does not induce any major dissociation of the oligomeric species present in solution (**Fig. 3B**). Independent 3D reconstruction performed on the smaller oligomeric species reveals a spheroidal particle similar in shape and size to the reconstructed model obtained in the absence of arsenous acid (**Fig. 3D**, middle panel).

**Figure 3.**
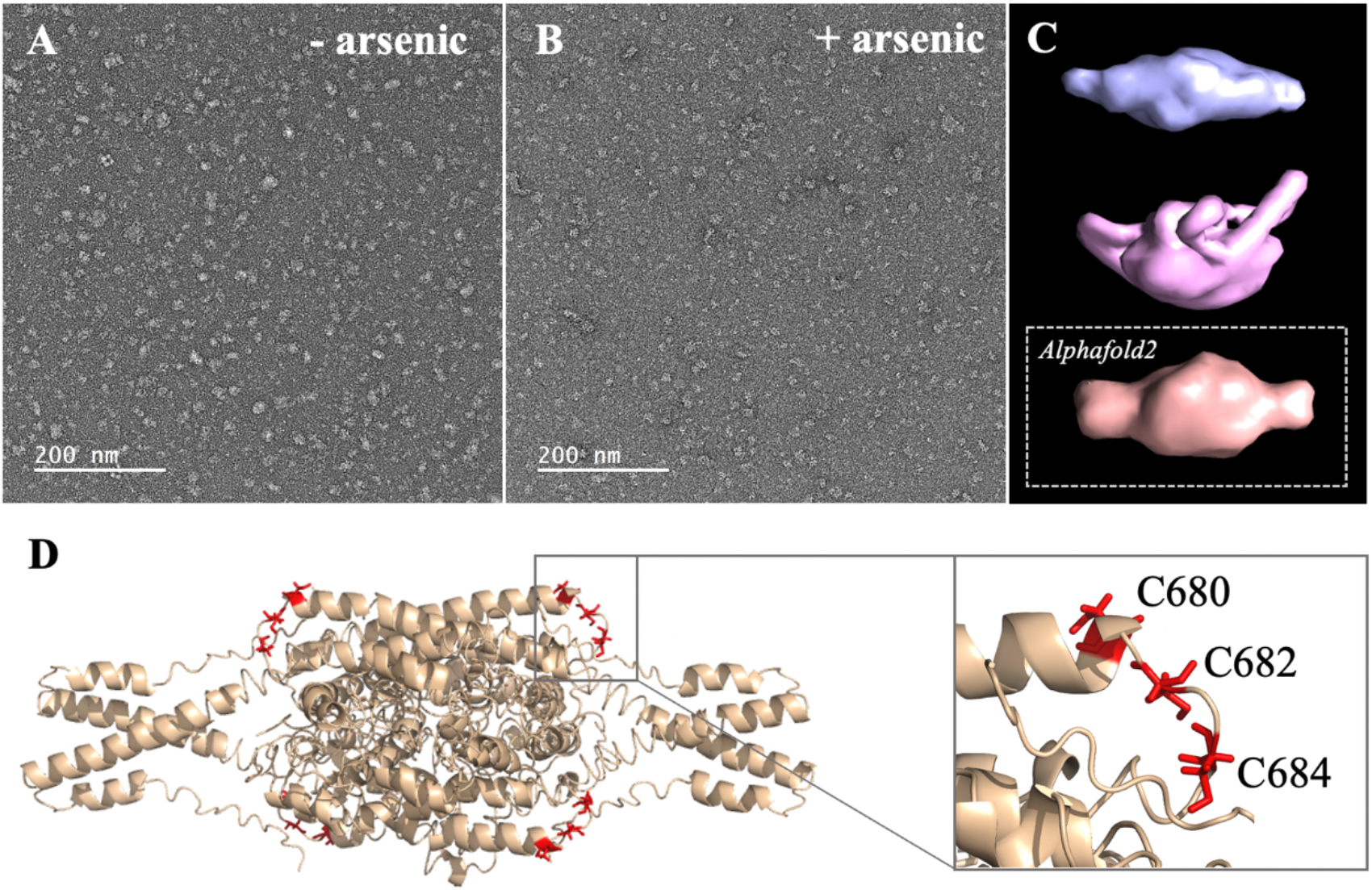
**(A-B)** Micrographs collected by TEM on a solution of the C-terminal WT construct in (**A**) absence of arsenous acid, and (**B**) in the presence of arsenous acid. (**C**) 3D models reconstructed from the smaller oligomeric species visible in the micrographs, with the upper panel (blue) showing the reconstructed model obtained in the absence of arsenous acid, the middle panel (pink) showing the model obtained in the presence arsenous panel, and the lower panel (orange) showing the envelop representation of the structural prediction obtained by Alphafold2. (**D**) Structural prediction of a tetrameric assembly of DISC1 C-terminal WT construct (WT) by Alphafold2, depicting the cysteine motif composed of C680, C682, and C684 in red. Insert shows that the cysteine motif is predicted to be located in a loop, fully exposed to solvent.

We then compare these low-resolution models with structural predictions from Alphafold2.^26^ We found that the best prediction for the tetrameric C-terminal DISC1 WT construct matches very well with the 3D reconstructed models calculated from the TEM experiments described above (**Fig. 3C**, lower panel). The tetrameric model predicted by Alphafold2 consists of a head-to-toe assembly of two dimeric units (**Fig. 3D**). Individual subunits are predominantly composed of coils and helix-turn-helix motifs, in good agreement with previous reports. The oligomerization interface is formed by a large helical bundle involving the C-terminal region of the domain (residues 689-834) while the N-terminal region forms a helix-turn-helix motif that is sticking out of the central core and gives the oligomer its spheroidal shape (**Fig. 3D**). Importantly, the cysteine motif encompassing C680, C682, and C683 is predicted to be located in a loop at the edge of the central bundle, fully exposed to solvent, and therefore readily accessible to arsenic (insert **Fig. 3D**). To complement the predictions by Alphafold2, we also performed a series of secondary structure analysis with RaptorX-SS8^27^, Jpred4^28^, and s2D^29^. All suggest that the cysteine motif is located within a loop predicted to be exposed to solvent.

The combination of SEC, TEM, and structural predictions, described above provide a molecular framework to explain the high affinity of DISC1 to arsenous acid. Altogether these results suggest indeed that the predominant species observed in our sample consist of a tetrameric complex of DISC1 C-terminal domain into which the arsenic-binding cysteine motif is fully exposed to solvent and readily accessible to arsenous acid.

## DISCUSSION

An ever-growing wealth of data has firmly established DISC1 as a multi-faceted scaffold protein that plays a pivotal role in mediating enzyme activity and protein-protein interaction for multiple signaling pathways involved in neurodevelopment, neural migration, and synaptogenesis. The role of DISC1 as a translational activator within Akt/mTOR pathway in particular has attracted a lot of attention over the past decade.^11,12^ Yet a precise analysis of structure-function relationship has proven to be extremely challenging. Structural characterization of DISC1 has long been hampered by its partially disordered nature and high-propensity for aggregation of recombinantly expressed constructs. In addition, the low degree of conservation of DISC1 sequence across species and lack of homology with other known proteins have prevented detailed identification of functional domains.^16,17^ In the present study, we examine the molecular basis of the recently reported ability of DISC1 to both detect and orchestrate the cellular response to oxidative stress induced by arsenic.^15^ The presence of a cysteine motif (C-X-C-X-C) in the C-terminal domain of DISC1 led us to investigate whether DISC1 could directly bind to arsenic. DISC1 C-terminal domain is especially relevant from a pathophysiological perspective because it encompasses polymorphisms recognized as risk factors for mental illness, such as the S704C and L807-frameshift mutant.^22,23^ The C-terminal domain is also deleted in the Scottish variant as a consequence of a balanced chromosomal translocation.^1,2^

Here, we designed a C-terminal construct (residues 598-854, **Fig. 1C**) encompassing the S (635-738) and C (684-836) structural regions identified by Korth and coworkers.^25^ Our C-terminal construct expresses recombinantly and predominantly forms tetramers in solution (**Fig. S1**). This matches well the observation by Korth and coworkers who reported that the S-region is predominantly tetrameric.^25^ Notably, the C-terminal contains a self-associating region rich in Ala, Gly, Leu, Pro, and Ser residues with high β-strand propensity, reminiscent of amyloid proteins. It has recently been reported that this region may promote the aggregation and/or fibrillization of DISC1 in the right physiological environment.^30^ Our purified samples presented no evidence of irreversible aggregation over time but it should be noted that the samples were kept at relatively low concentration (2 μM) in conditions never exceeding room temperature. Nevertheless, the TEM micrographs clearly show that even after elution through SEC, a significant fraction of the samples form large amorphous aggregates, indicating the C-terminal domain has indeed a high-propensity to self-associate beyond tetrameric states (**Fig 3A-B**).

Our C-terminal construct was used in fluorescence-based assays to determine the affinity to trivalent arsenous acid, which is known to bind to the thiol groups of cysteines. We found that the WT variant (598-854) that contains a total 7 cysteines (i.e. three cysteines composing the C-X-C-X-C motif: C680, C682, and C684, and four isolated cysteines: C698, C762, C782, and C844), shows clear evidence of binding in the low micromolar range (**Fig. 2**). We then compared the WT construct with a pseudo-WT variant named WT* where the four isolated cysteines (C698, C762, C782, and C844) were mutated to serine (**Table 1**). This WT* variant presented a similarly high affinity toward arsenous acid compared to the reference WT construct, although with a slightly lower maximum binding capacity (**Fig. S2A, Table 1**), indicating that arsenous acid is specifically binding to the three-cysteine motif (C680, C682, C684). Next, we used a series of single, double, and triple mutants to determine whether all three cysteines composing the motif were required to stably bind arsenous acid. Fluorescence assays recorded for the single mutants (C680S*, C682S*, and C684S*) reveal a large decrease in binding capacity compared to the WT or WT* variants (**Fig. S2, Fig S2B**, and **Table 1**), indicating all three cysteine of the motif participate in stabilizing interaction with arsenous acid. Notably, the consequence of the mutation was less important for C680 (**Table 1**), which suggests a lower contribution of the first cysteine of the motif toward binding. The structural model provided by TEM and Alphafold2 shows indeed that C680 is located at the C-terminal tip of an α-helix while C682 and C684 are positioned in a loop fully exposed to solvent and likely more readily available to bind arsenous acid (**Fig. 3D**). Analysis of the binding assays recorded for the double and triple mutants shows binding capacities within the same range of those measured for the single mutants (**Fig. 2, Fig. S2C**, and **Table 1**), which confirms that all three cysteines of the motif participate to some degree in arsenous acid binding.

Next, we collected a series of micrographs by TEM of DISC1 C-terminal domain, in the absence and in the presence of arsenous acid, to gain further insights into the molecular basis of DISC1 ability to bind arsenic (**Fig. 3 A-B**). 3D reconstruction performed on the smaller oligomeric species observed in the micrographs reveals spheroid complexes compatible with a tetrameric complex (**Fig. 3C**). 3D reconstructed models obtained in the absence and in the presence of arsenous acid were found to be virtually identical indicating that binding to arsenous acid does not induce dissociation or major structural reorganization of the tetrameric complexes (**Fig. 3C**). Finally, a series of structural predictions were run to complement the results obtained by TEM. Predictions by Alphafold2 for a tetrameric complex of the C-terminal domain of DISC1 matches very well the size and dimensions of the 3D reconstructed models (**Fig. 3C**). The structural details of the prediction suggest a head-to-toe assembly of two dimeric units held together by a central helical bundle. The oligomeric interface consists of a series of helix-turn-helix motifs involving residues from the C-terminal region of the domain. This is in good agreement with recent report by Cukkemane et al. describing this region (residues 717-761) as a self-association core.^30^ Prediction by Alphafold2 in terms of local secondary structure also matches well data reported on truncated C-terminal fragments in complex with a camel nanobody and peptide fragments of ATF4 and NDE1.^19-21^ Importantly, these predictions suggest that the cysteine motif is located in a loop fully exposed to solvent (**Fig. 3D**), which provides a clear molecular framework explaining the ability of DISC1 to bind arsenous acid.

Altogether, these results provide insights into yet another functional facet of DISC1. As an arsenic binding protein, DISC1 could potentially act both as a cellular sensor and direct translational activator in response to the resulting oxidative stress. Our TEM data show no major conformational change of the C-terminal domain in the presence of arsenic but due to the rather low spatial resolution of this technique, one can’t exclude that arsenic may induce a subtle rearrangement of local structural motifs. Furthermore, in a cellular environment, arsenic may affect interactions between DISC1 and proteins partners such as girdin, which could lead to a switch of DISC1 function from sensor to translational activator (**Fig. 1 A-B**).

## EXPERIMENTAL PROCEDURES

### Plasmid design and mutagenesis

DNA sequence encoding DISC1 WT construct (amino acids 598-854) and other variants (**Table 1**) were inserted into the pET28b (+) vector as His-tag fusion protein. All carry the additional mutations W602F and W752F, leaving a single tryptophan (W691) as a reporter for the fluorescence assays. These mutations were generated by QuickChange site directed mutagenesis and confirmed by DNA sequencing (Iowa State University DNA Sequencing Facility).

### Protein Purification

All constructs were transformed into E. coli strain BL21 (DE3). Cells were grown in LB media at 37 °C with kanamycin was added to a final concentration of (50 μg/mL). Growth was monitored by absorbance at wavelength 600 nm. At A600 of 0.7, the temperature was reduced to 25 °C and IPTG was added to a final concentration of 0.25 mM. Cells were collected after 16-20 hours of induction and suspended in a lysis buffer (Table 2). DNase (1 mg/mL), leupeptin (1 mg/mL), and PMSF (0.1 M) were all added to the solution. Cells were lysed by sonication and then centrifuged at 16000 rpm for 1 hour. Lysate was loaded into a Ni-NTA column equilibrated with W1 buffer (50 mM Tris-HCl, pH 8.0, 5 mM NaCl, 5 mM Imidazole). Ni-NTA column was washed with 300 mL of W1 buffer and 150 mL of W2 buffer (50 mM Tris-HCl, pH 8.0, 5 mM NaCl, 30 mM Imidazole) and 200 mL of salt buffer (50 mM Tris-HCl, pH 8.0, 400 mM NaCl, 30 mM Imidazole). Samples were eluted with elution buffer (50 mM Tris-HCl, pH 8.0, 5 mM NaCl, 250 mM Imidazole) then dialyzed overnight against 4 L of dialysis buffer (50 mM Tris-HCl, pH 8.0, 5 mM NaCl). The samples were then loaded into the Size Exclusion Chromatography (SEC) HiLoad® 26/600 Superdex® 200 pg pre-equilibrated with running buffer (50mM of Tris-HCl, 10mM of NaCl at pH 8.0) for further purification. Finally, purified proteins were examined by SDS-PAGE (15%, w/v) to ensure purity of at least 85%.

### Tryptophan fluorescence assays

Fluorescence binding assays were performed using an excitation wavelength of 284 nm and emission wavelength of 348 nm. Ultraviolet (UV) light was applied as the light source with a PMT of 800. Arsenous acid stock solution was purchased from Thermo Fisher Scientific and diluted in fluorescence assay buffer (50mM Tris-HCl, 10mM NaCl at pH 8.0) and stored at room temperature. Protein samples were added to a final concentration of 2 μM in 2 mL the quartz cuvettes with mini stir bar. A blank buffer reading was recorded as baseline and subtracted to the raw fluorescence signal. Each protein sample was titrated in triplicate to estimate average and standard deviations. Titration profiles were collectively fitted (i.e. including all variants presented in Table 1) to a two-site specific binding model in Graphpad Prism 8 to estimate the dissociation constants, K_d_ (high) and K_d_ (low), and maximum binding capacity.

### Transmission electron microscopy (TEM) measurements

The sample solution (4 μl) containing 10 μM protein was applied onto a glow-discharged carbon-coated copper grid (S160-4, Plano). After 2 min, the solution on the grid was blotted off by filter paper. The grid was then washed with 4 μl of 1% (w/v) uranyl acetate (UrAc) and blotted off immediately, and another 4 μl of 1% (w/v) UrAc was applied onto the grid for 1 min. TEM images were obtained using a TFS Talos 120 C (Thermo Scientific) with a voltage of 120 kV. Processing of the negatively stained images was performed using RELION 3.1.^31^ Contrast transfer function (CTF) was fitted using CTFFIND4.^32^ For the tetrameric oligomers, 40 particles were selected from ten images (pixel size of 2.5 Å) and extracted with a box size of 80 pixels (200 Å). 2D classification was performed with a mask diameter of 180 Å.

### Structural predictions

Secondary prediction was performed using RaptorX-SS8^27^, Jpred^28^, and s2D^29^ using the sequence of the human DISC1 C-terminal domain (Uniprot Q9NRI5-1, residue 598-854) as template. Complete structural prediction of DISC1 598-854 was obtained for the DISC1 C-terminal domain using Alphafold2^26^ provided by the High-Performance Computing facility at Iowa State University.

## Supporting Information

This article contains supporting information

## Funding and Additional Information

This project is supported by funds from the Roy J. Carver Charitable Trust of Muscatine, Iowa, and from NIGMS R01 GM132561 (J.R). The content is solely the responsibility of the authors and does not necessarily represent the official views of the National Institutes of Health.

## Acknowledgements

We acknowledge the Roy J. Carver High Resolution Microscopy Facility (Office of Biotechnology, Iowa State University, Ames, IA) for providing analytical instrumentation and we are thankful to Tracey P. Stewart for helping with TEM set up and data collection.

