## Supplementary material for "Evidence of Disrupted-in Schizophrenia 1 (DISC1) as an arsenic binding protein and implications regarding its role as a translational activator": Fig. S1, Fig. S2

### SUPPLEMENTARY FIGURES

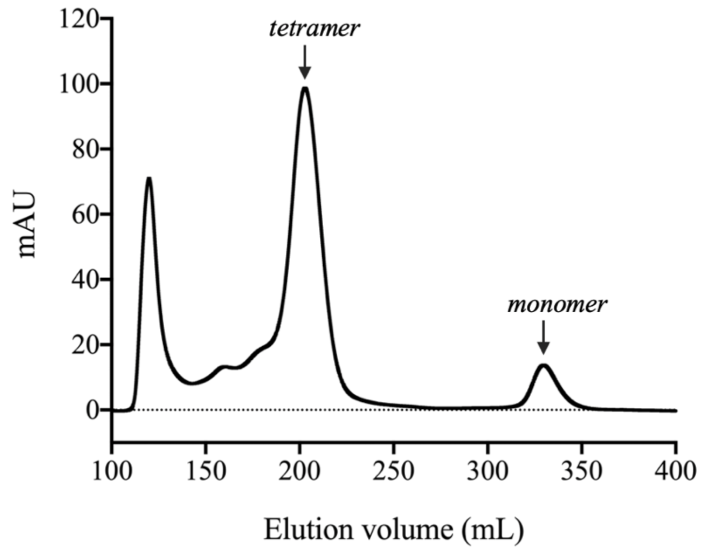


**Figure S1.** Size-exclusion chromatography elution profile obtained for the DISC1 C-terminal construct (WT) highlighting the peaks corresponding to monomeric and tetrameric species. The peak eluting between 100-150 mL contains high molecular weight oligomers and amorphous aggregates.

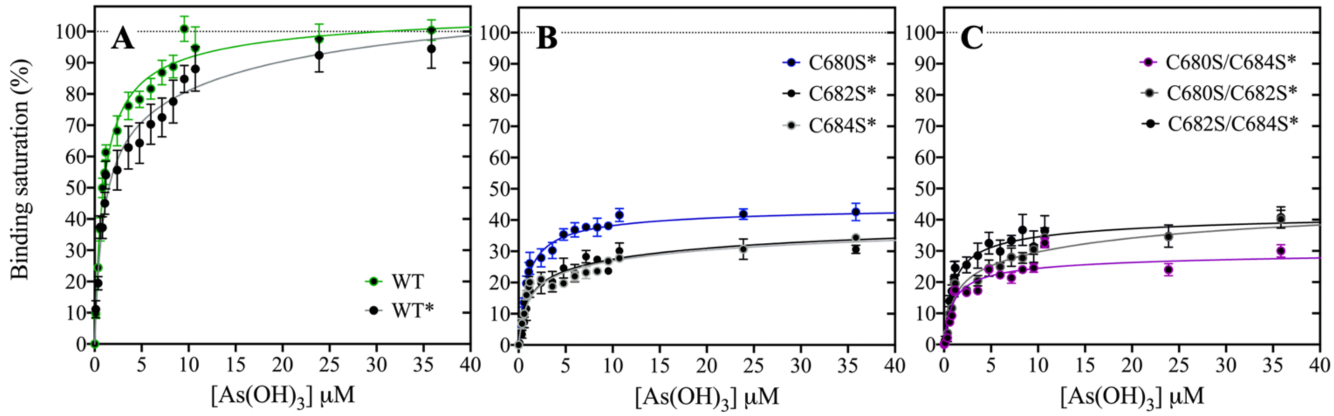


**Figure S2.** Plots of depicting the binding saturation as a function of arsenous acid concentration obtained for (**A**) WT and WT*, (**B**) the single cysteine variants C680S*, C682S*, and C684S*, and (**C**) the double mutants C680S/C682S*, C680S/C684S*, and C682S/C684S*. Lines correspond to a collective fit of all data presented in Table 1 using a simple two-site specific binding model.
